# Stability of brain-behavior correlation patterns across measures of social support

**DOI:** 10.1101/2023.03.23.533966

**Authors:** Haily Merritt, Joshua Faskowitz, Marlen Z. Gonzalez, Richard F. Betzel

## Abstract

The social environment has a critical influence on human development, cognition, and health. By using network approaches to map and analyze the connectivity between all pairs of brain regions simultaneously, we can clarify how relationships between brain regions (e.g. connectivity) change as a function of social relationships. Here we apply multilayer modeling and modularity maximization–both established tools in network neuroscience–to jointly cluster patterns of brain-behavior associations for seven social support measures. Our analyses build on both neuroecological findings and network neuroscientific approaches. In particular we find that subcortical and control systems are especially sensitive to different constructs of perceived social support. Network nodes in these systems are highly flexible; their community affiliations, which reflect groups of nodes with similar patterns of brain behavior associations, differ across social support measures. The multilayer approach used here enables direct comparison of the roles of all regions of the brain across all social support measures included. Additionally, our application of multilayer modeling to patterns of brain-behavior correlations, as opposed to just functional connectivity, represents an innovation in how multilayer models are used in. More than that, it offers a generalizable technique for studying the stability brain-behavior correlations.

## Introduction

Research in health psychology and social neuroscience indicate an urgent need to understand how social support is related to whole-brain connectivity (Xiao et al., 2021). This endeavor involves linking many strands of research; to date, studies have connected perceived social support to wellbeing (Cohen, 2004; Williams et al., 2018), experience in the environment to brain network organization (Chan et al., 2018; Ellwood-Lowe et al., 2021), and the social world to the brain (Krendl and Betzel, 2022; Schmälzle et al., 2017; Morawetz et al., 2021; Rudolph et al., 2021).

More than an arena for social cognition to play out, the social environment is home to resources important for human cognition (Gross and Medina-DeVilliers, 2020) and wellbeing (De Jaegher et al., 2010; Coan et al., 2017; Morawetz et al., 2021; Zaki and Williams, 2013). The quality of our social relationships and the support we receive from them has a large impact on our health (Beckes and Sbarra, 2022), including better prognoses related to disease (Cohen and Wills, 1985; Cohen, 1988, 2004; Williams et al., 2018), increased antibody response to vaccines (Uchino et al., 2020), lower levels of inflammation (Uchino et al., 2018), reduced physiological impact of adversity (Wymbs et al., 2020), conserved energy in response to threats (Gonzalez et al., 2021; Saxbe et al., 2020), decreased risk of mortality (Berkman and Syme, 1979), and more (Miller et al., 2009; Seeman and McEwen, 1996; House et al., 1988; Uchino et al., 1996; Hawkley and Cacioppo, 2007; Repetti et al., 2002; Chen and Miller, 2007).

From a neuroecological perspective, social support can be seen as a resource, reliable access to which can dramatically alter health and behavioral outcomes. Indeed, neuroecological approaches have long investigated the link between social relationships and brain connectivity Gonzalez et al. (2015); Coan et al. (2017); DeCross et al. (2022); Morawetz et al. (2021); McLaughlin et al. (2014); Sheridan and McLaughlin (2014); Gee et al. (2013); Callaghan and Tottenham (2016). While neuroecologists, who study how brain organization and dynamics vary across individuals given the experiences during development, often have extensive data about individuals’ environments, brain analyses from a neuroecological perspective often focus on the connectivity between a few seed regions of interest (Gonzalez et al., 2015; Gee et al., 2013; Callaghan and Tottenham, 2016, but see Pelletier-Baldelli et al., 2022; DeCross et al., 2022 for network approaches). Network neuroscience, on the other hand, provides tools for understanding environmentally-influenced differential connectivity in the context of the entire brain. Although we are far from a thorough understanding of brain network reconfigurations in response to dynamic social environmental resources, integrating network and neuroecological perspectives serves to advance both fields.

Network techniques take advantage of whole-brain data by considering all interactions between all regions of the brain simultaneously. This holistic approach clarifies the contributions of regions of interest in the context of the whole brain. Importantly, whole-brain network approaches do not preclude using extensive data about the environment; indeed, in recent years many studies have made connections between changes in brain networks over the course of development (Uddin et al., 2011; Fair et al., 2007; Dosenbach et al., 2010; Betzel et al., 2014; Baum et al., 2017) and variation in brain networks across different environmental experiences (Chan et al., 2018; Ellwood-Lowe et al., 2021; Chen et al., 2016; Rakesh et al., 2021).

Moreover, a variety of network tools (e.g. multilayer networks) exist to study brain connectivity over time or across contexts. When network neuroscientists have considered social information, the focus is often on social cognition and affect (Barrett and Satpute, 2013), social networks (Noonan et al., 2018), face perception (Cassidy et al., 2021), emotion regulation (Morawetz et al., 2021, 2020), loneliness (Mwilambwe-Tshilobo et al., 2019; Spreng et al., 2020), or socioeconomic status (Chan et al., 2018; Ellwood-Lowe et al., 2021). Understudied this work, then, is a network neuroscientific approach to *social support*. How do whole-brain patterns of connectivity vary across different experiences of perceived social support? Addressing such a question would contribute to ongoing work linking peoples’ *subjective* perceptions to neural data, like fMRI (Mwilambwe-Tshilobo et al., 2019; Spreng et al., 2020), which, for social information, can yield different and additional insights than objective measures of social experience alone (Coan et al., 2017; Chen et al., 2004).

The present study builds on existing work linking the social environment to variation in brain network organization. Specifically, we use a well-established multilayer modeling framework (Mucha et al., 2010) to examine correlations of brain functional connectivity weights with a suite of social support measures. In a multilayer model, each layer is an individual network (e.g. a single functional connectivity network) which can be coupled to other layers to measure change in a network across the layers. Whereas past research has used the same framework to track variation in network structure across time (Bassett et al., 2011; Betzel et al., 2017; Finc et al., 2020), individuals (Betzel et al., 2019), or connectivity modalities (Puxeddu et al., 2022), we use this technique to track variation in brain-behavior correlation patterns. Our application of the multilayer framework presents a series of conceptual advances: (1) it preserves correlation profiles of individual social support measures, conferring a greater degree of interpretability, and (2) it identifies groups of brain regions whose correlation profiles are stable *versus* variable across social support measures, i.e. inflexible *versus* flexible, providing a substantive contribution to the science of neural sensitivity to dimensions of social context. Put simply, we link the intricacies of social environment experience to the intricacies of brain network connectivity, without dimensionality reduction on either data source to make the connection

## Results

Our data included self-report social support data and resting state fMRI (rsfMRI) data from 100 unrelated adult subjects from the Human Connectome Project (HCP) (Van Essen et al., 2013). For the social data, we used the seven perceived social support measures from the NIH Toolbox included in HCP data: *Friendship, Emotional Support, Instrumental Support, Hostility, Rejection, Loneliness, and Stress* (Cyranowski et al., 2013). For the brain data, we used four resting state functional magnetic resonance imaging (fMRI) scans. The social and brain data were collected within one day of each other.

### Permutation test of correlations between edge weights and perceived social support

To test which correlations between edge weights and social support measures had a greater magnitude than would be expected by chance, we used permutation testing. We shuffled social support scores across subjects (i.e., we shuffled the order of the rows of the matrix in Figure 1b) 1000 times and recalculated the correlation between edge weights and social support scores each time. We counted the number of instances for which the observed correlation between edge weight and social support score had a greater magnitude than the permuted correlation. To obtain a *p*-value for each edge weight-social support measure correlation, we divided this count by the number of permutations (1000) and used false-discovery rate of 0.05 to correct for multiple comparisons. This gave an adjusted significance criterion of *p*_*crit*_ *<* 0.046. Figure 2 shows which elements and what proportion of each canonical system survived permutation testing.

**Figure 1:**
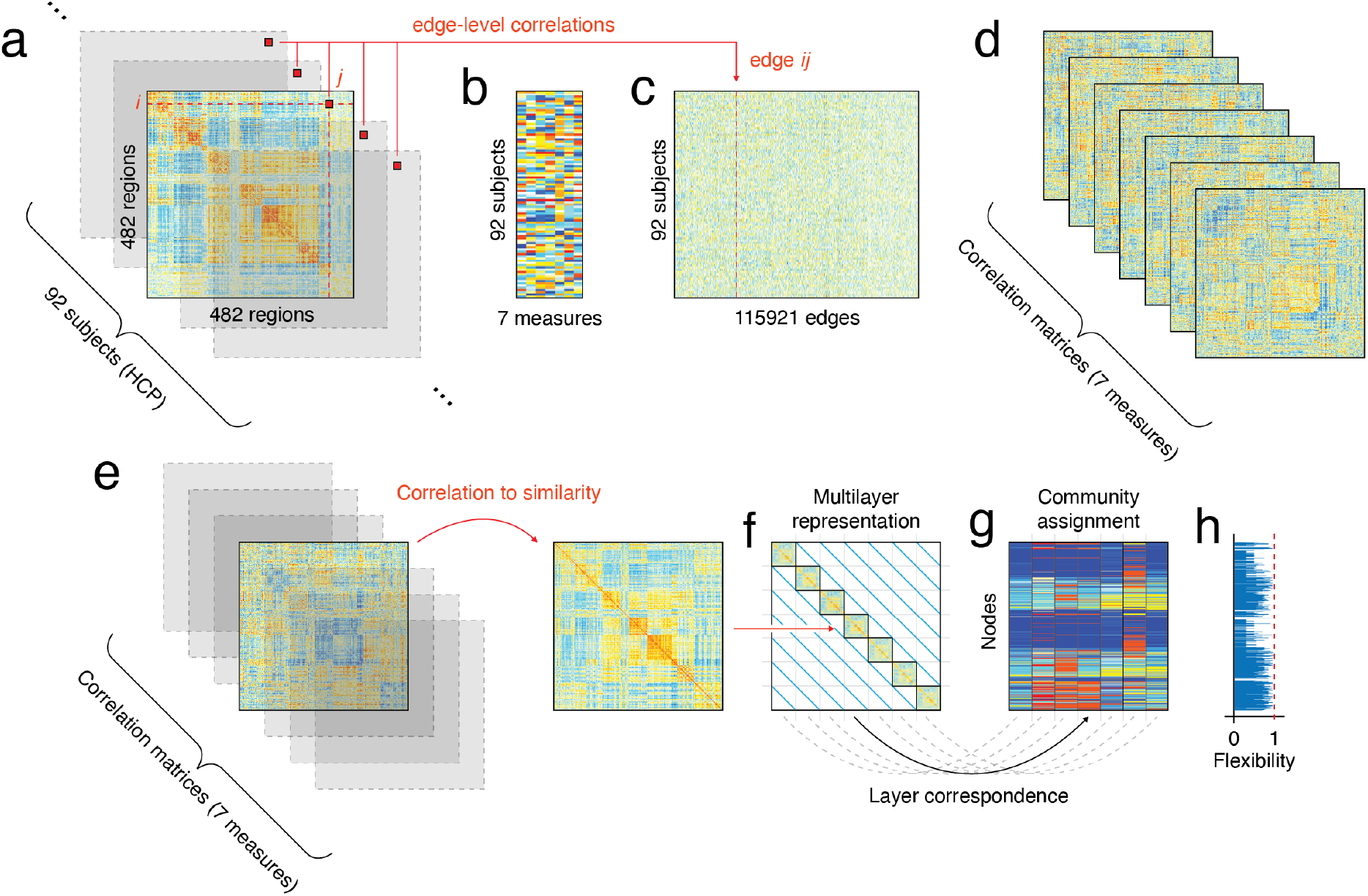
A schematic of the method for computing the correlations between functional connectivity and social support. **(a)** shows the 92 (one per subject) functional connectivity matrices, where element *ij* indicates the Pearson correlation between nodes *i* and *j*. (b) represents all 92 subjects scores on seven multi-item measures of perceived social support: *Friendship, Stress, Rejection, Hostility, Emotional Support*, and *Instrumental Support*. The matrices in (a) are vectorized and stacked on top of each other in (b) to yield a 92 (number of subjects) by 115921 (number of edges on each functional connectivity matrix) matrix. We compute the correlation between social support measures (b) and edge weights (c), resulting in seven (one for each measure) correlation matrices (d), where element *ij* in matrix *k* gives the correlation between social support measure *k* and the edge weight between nodes *i* and *j*. Because modularity maximization treats negative weights as belonging to just one community, we transform each correlation matrix to a similarity matrix (e). We do this by computing the similarity between every pair of rows in the correlation matrix. Each of the seven similarity matrices becomes itself a layer in the multilayer model (f). The multilayer model is a 2D tensor (f) whose on-diagonal elements are the seven correlation matrices and whose offdiagonal elements are the coupling between layers. We perform community detection on this multilayer model, which yields a community assignment for every node for each layer (g). We compute for each node its flexibility, or how much it changes its community affiliation between layers (h).

**Figure 2:**
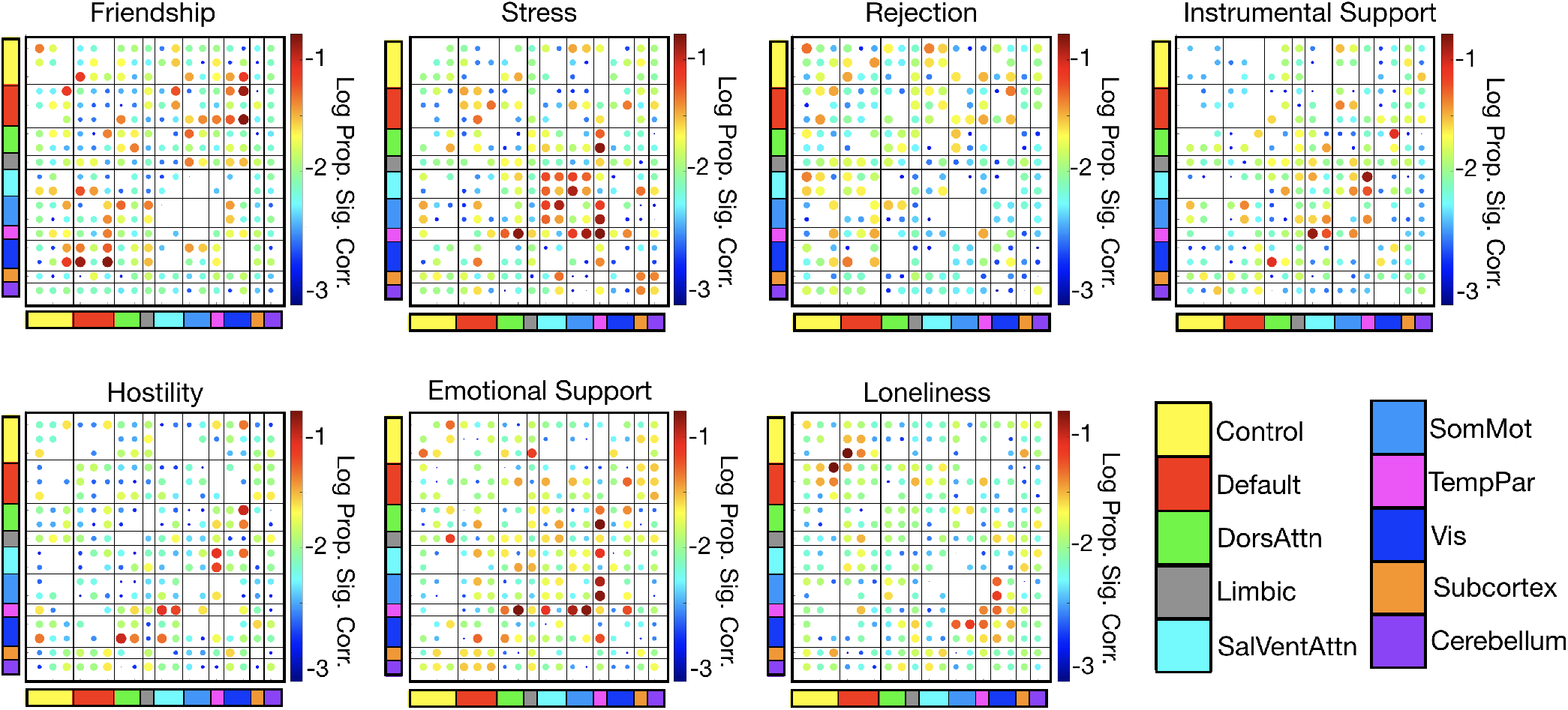
System-level correlations between edge weights and social support measures. We shuffled social support scores across subjects and recalculated the correlations with edge weights 1000 times. For each measure, we summarize the results of this permutation testing by showing the log proportions of edges of each canonical subsystem (e.g. ContA, ContB) that are significantly correlated with the given social support measure using a false discovery rate of 0.05 (adjusted *p*_*crit*_ = 0.046).

### Analyses of the parameter space of the multilayer model

How do system- and layer-specific behaviors change as community resolution *γ* and inter-layer coupling *ω* change? To answer this, we analyzed the parameter space of our multilayer model using several methods: applying principal components analysis to measure of community flexibility to identify modes of community variation across parameter values, by tracking variability in regional flexibility patterns, by studying the partition landscape degeneracy (a measure of how similar detected partitions are to one another at given point in parameter space), and by comparing select partition “seeds” to the rest of the parameter space. These analyses motivate our selection of points in parameter space for further study.

#### Principal components analysis

First, with the aim of arriving at a holistic understanding of how varying the parameter values is related to changes in the relationship between brain networks and social support, we performed a principal components analysis (PCA) on the flexibility of all nodes in all layers across all values in parameter space (see S2 for amount of variance explained) Betzel et al. (2019). Figure 3a shows the magnitude of expression of the first principal component, which explains about 35% of the variance, by each point in parameter space. Our results suggest that strong positive coefficients were associated with small values of the community resolution parameter *γ*, and strong negative coefficients with larger *γ*s. Figure 3b projects this into brain space for both the cortex and subcortex. The point in parameter space whose coefficient for principal component 1 had the largest magnitude was at *γ* = 0.15, *ω* = 0.001. At these parameter values, there were 14 communities with an average size of 241 *±* 109.37 nodes.

**Figure 3:**
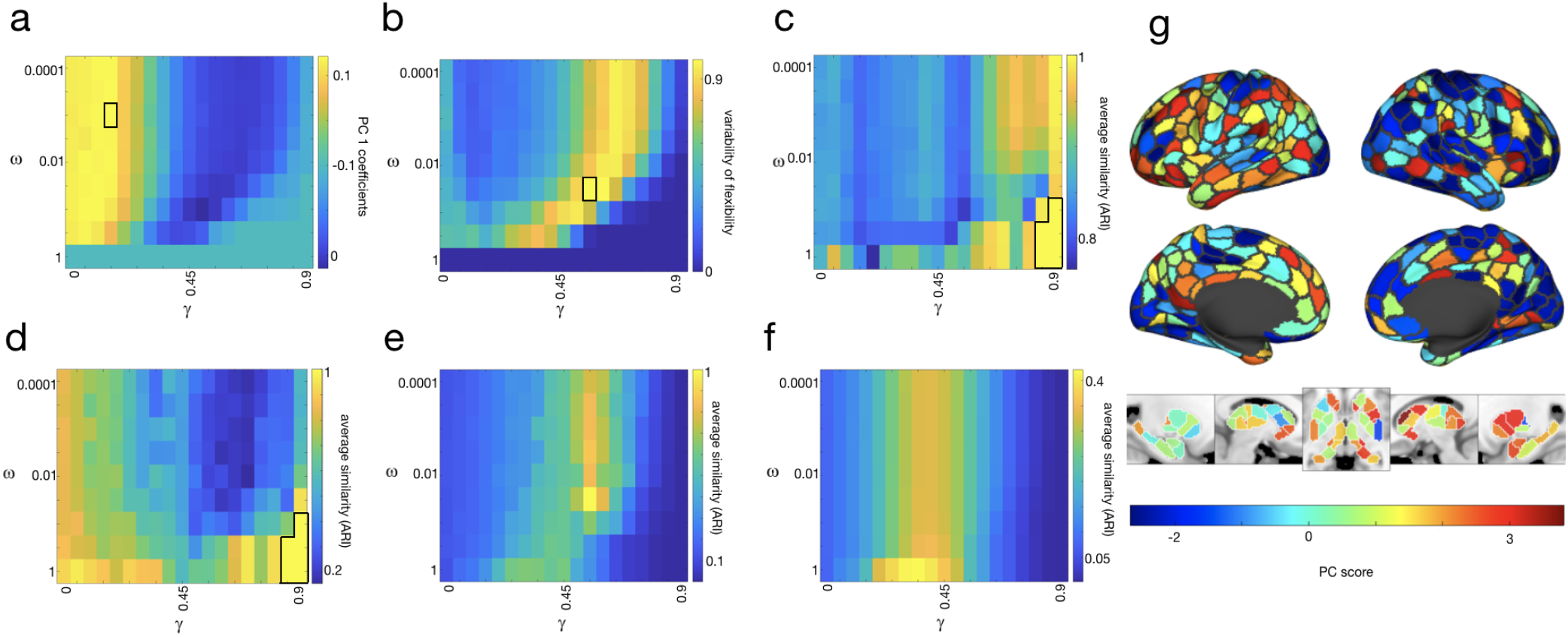
Analyses of parameter space. We performed PCA on the entire parameter space; (a) shows the magnitude of expression of the first principal component. The point in parameter space for which the magnitude of expression of the first principal component is greatest is *γ* = 0.15, *ω* = 0.001, indicated by a black outline. (b) shows the variability in node flexibility across parameter space computed across individual partitions at a given point in parameter space. The point in parameter space that maximizes variability in flexibility is *γ* = 0.55, *ω* = 0.0316, indicated by a black outline. In (c), for every point in parameter space, the similarity between every partition was computed using the adjusted Rand Index then averaged for each point across layers. For this calculation, the global maximum similarity is achieved by five points in parameter space: *γ* = 0.9, *ω* = 0.1; *γ* = 0.85, *ω* = 0.3162; *γ* = 0.9, *ω* = 0.3162; *γ* = 0.85, *ω* = 1 and *γ* = 0.9, *ω* = 1, indicated by a black outline. We also calculated the similarity of each individual partition to the consensus communities at that point in parameter space and took the average across partitions and layers (d). For this calculation, the global maximum was reached by a set of five points in parameter space: *γ* = 0.9, *ω* = 0.1; *γ* = 0.85, *ω* = 0.3162; *γ* = 0.9, *ω* = 0.3162; *γ* = 0.85, *ω* = 1 and *γ* = 0.9, *ω* = 1, indicated by a black outline. Panels (e) and (f) show seed-based analyses, where we computed the similarity of all partitions for each point in parameter space to the seed. For (e), we used as a seed *γ* = 0.55, *ω* = 0.0316, which maximizes the variability in flexibility. For (f), we used as a seed *γ* = 0.35, *ω* = 1, which is the point in parameter space whose communities are most similar to the Yeo systems. (g) shows the extent to which different regions of the cortex and subcortex express the mode of flexibility present in the first principal component.

#### Variability in flexibility

Second, we computed for each point in parameter space the variability of flexibility for all nodes across all layers (see Figure 3c). Previous work (Bassett et al., 2011) has selected the parameter values which maximize variability in flexibility for further analyses. In our case, this point was at *γ* = 0.55, *ω* = 0.0316. With this pair of parameter values, we detected 79 communities with an average size of 42.71 *±* 10.67 nodes.

#### Partition landscape degeneracy

Given that we ran 100 iterations of community detection, how similar was each resulting partition to other runs with the same parameter values? That is, for which points in parameter space did the community detection algorithm consistently find the same communities across runs? We approached this two ways. First, for a given pair of parameter values, we calculated the similarity (using the adjusted Rand Index; ARI) between every pair of partitions then took the average (see Figure 3d). In general, a larger ARI for a pair of partitions indicates that they are relatively similar to each other; smaller values mean greater dissimilarity between partitions. Second, for every pair of parameter values, we calculated the similarity of each partition to the consensus communities (see Figure 3e). This second approach tells us more about how representative the consensus communities are for a given pair of parameter values. For both of these approaches, similarity was maximized at the highest values of the community resolution parameter *γ* (i.e., many small but internally very similar communities) and the coupling parameter *ω* (i.e., maximal inter-layer coupling).

#### Seed-based similarity

Next, we wanted to quantify how similar a community partition at one point in parameter space was to all other points. In other words, how representative of all of parameter space is the one pair of *γ* and *ω* values that maximize variability in flexibility? We approached this two ways. First, we selected the point in parameter space whose consensus communities maximized variability in flexibility (*γ* = 0.55, *ω* = 0.03). Then, we used this point as a seed and computed the adjusted Rand Index to the seed for the consensus communities for all other points in parameter space (see Figure 3e). Additionally, to link our results more clearly to canonical brain systems in network neuroscience, we also used as a seed the point in parameter space that maximized similarity (using ARI) to the Yeo systems (Yeo et al., 2011). This point was found at *γ* = 0.35, *ω* = 1, when layers are maximally coupled (see Figure 3f).

#### Selecting points in parameter space for further analysis

In principle, any of these approaches could be used to select a point or points in parameter space for further analysis. In general, we saw more variation across values of the community resolution parameter *γ* than the coupling parameter *ω*, in line with other studies (Betzel et al., 2019). Coefficients for the first principal component suggested different patterns are to be found for smaller *γ*s versus larger *γ*s. To select specific smaller and larger *γ* values, we opted for the two points in parameter space whose coefficients on the first and second principal components have the greatest magnitude. The parameter values for which the magnitude of the coefficient of the first principal component is maximized is *γ* = 0.15, *ω* = 0.001. For principal component two these values are *γ* = 0.35, *ω* = 0.01. By using this approach, we have identified points are maximally representative of the dominant modes of variation in parameter space.

### Multilayer community structure

What does the multilayer community structure look like at the points in parameter space we have selected? When the magnitude of the coefficient for the first principal component is maximized (*γ* = 0.15, *ω* = 0.001), we find a total of 14 communities, only four of which were present in every layer (see Figure 4). Across the seven layers of perceived social support, a few patterns were consistent. Somatomotor and adjacent nodes tended to be in the same community in dark blue. For most layers, visual nodes were also in this same community. Frontal, temporal, medial, and subcortical nodes had more variety across layers.

**Figure 4:**
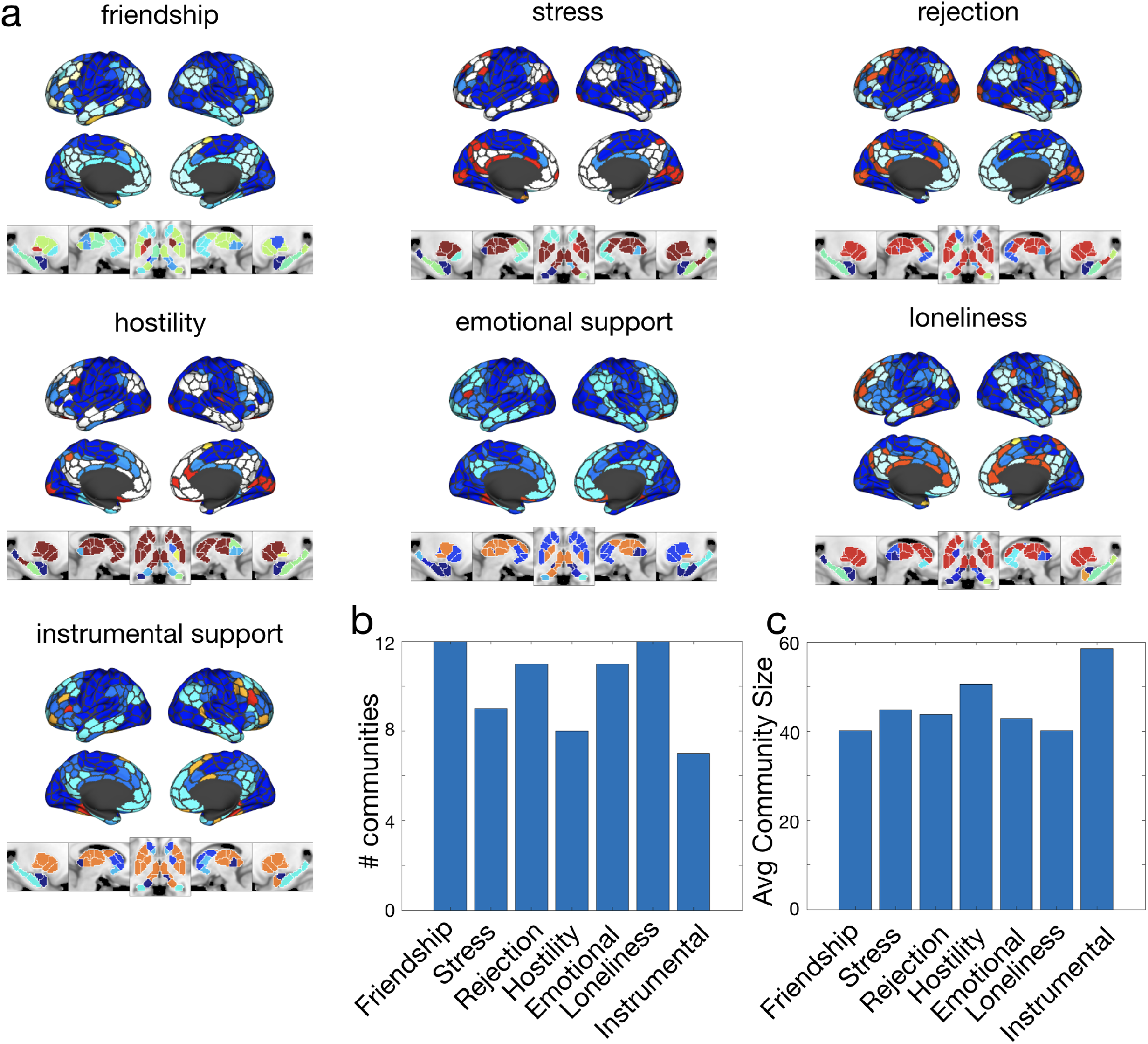
Multilayer consensus community structure. Panel (a) shows the consensus community structure across social support measures at the point in parameter space whose coefficient for principal component 1 has the largest magnitude (*γ* = 0.15, *ω* = 0.001). Colors indicate communities. There are 14 communities total, but only four communities are present across all layers (shown in dark blue, light blue, teal, and red). Panel (b) shows the number of communities in each layer according to the consensus partition. Panel (c) gives the average community size for each layer according to the consensus partition. The MNI coordinates for the five panels of subcortical communities are, from left to right: x = -23, x = 10, z = -3, x = 13, x = 27.

When the magnitude of the coefficient of the second principal coefficient is maximized (*γ* = 0.35, *ω* = 0.01), there were 25 communities, all of which were present in every layer (see Supplementary Figure S5). Somatomotor and visual nodes again tended to be in the same community, but with less consistency. While there were many communities represented by the subcortex, a single community (yellow) was most prominent across all layers, unlike when *γ* = 0.15, *ω* = 0.001. Again, frontal, temporal, and medial nodes were associated with a variety of communities across layers. To understand in more depth which nodes change their community affiliation between which layers, we looked at flexibility.

### Flexibility

#### Flexibility across the brain and across canonical systems

Which regions of the brain switch their community affiliation more (i.e., are more flexible)? Do flexible nodes tend to belong to the same canonical system? To address these questions, we examined the flexibility (i.e., the amount a node switches communities between layers) of nodes across the brain, across canonical systems, and across social support layers. We found that, at the point in parameter space where the magnitude of the coefficient for the first principal component is maximized (*γ* = 0.15, *ω* = 0.001), flexibility was widely distributed across the brain (see Figure 5a) but unevenly distributed across the nodes of canonical systems (*χ*^2^ = 48.46, *df* = 9, *p <* 0.0001 using a Kruskal-Wallis rank-sum test; see Figure 5b). The system with the most flexible nodes was ContC (median flexibility = 0.849), and the system with the least flexible nodes was SomMotA (median flexibility = 0.63).

**Figure 5:**
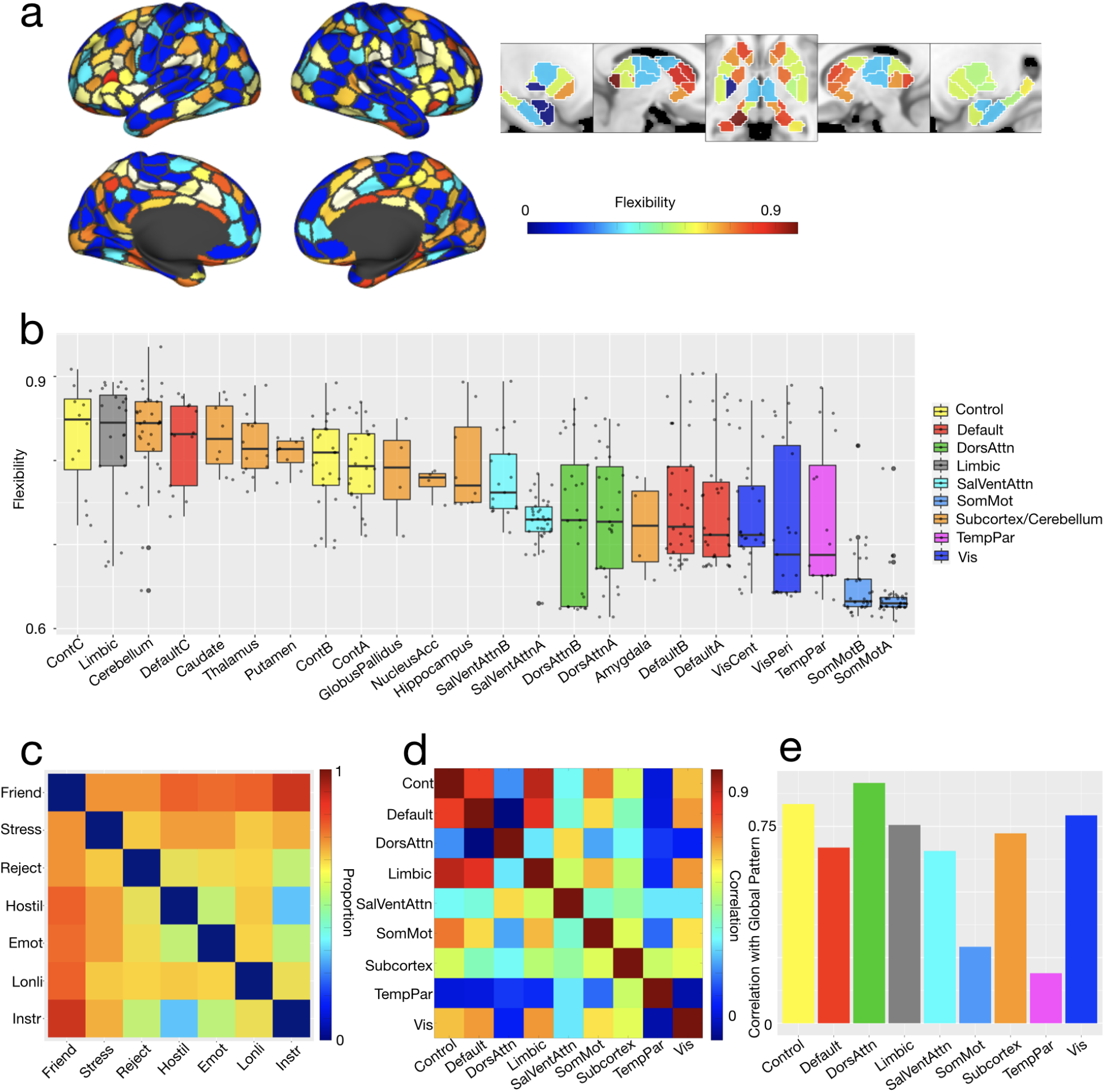
Flexibility. Panels (a), (b), (c), and (d) represent data from the point in parameter space (*γ* = 0.15, *ω* = 0.001) with the largest loading on the first principal component. Panel (a) shows the flexibility of cortical and subcortical regions. Panel (b) shows the distribution of flexibility across canonical systems, ordered by median flexibility. Panel (c) shows the proportion of all nodes that change community affiliation between each pair of the seven layers: Friendship, Stress, Rejection, Hostility, Emotional Support, Loneliness, and Instrumental Support. Panel (d) shows how similar all systems are to each other in terms of the proportion of nodes changing community between each pair of layers. Panel (e) plots the correlation between each system and the global pattern of community change between layers. The MNI coordinates for the five panels of subcortical flexibility are, from left to right: x = -23, x = 10, z = -3, x = 13, x = 27.

To compare the flexibility of nodes in different systems, we used permutation testing to construct a null model. We permuted all nodes’ flexibility 10000 times; for each permutation we compared the flexibility of the nodes for all pairs of systems. To compute *p*-values, we counted the number of instances in which the direction of difference in flexibility between a pair of systems was greater in the observed data than in the permuted data, then divided this by the number of permutations (10000). ContC emerged as more flexible than DefaultA, DefaultB, DorsAttnA, DorsAttnB, SomMotA, SomMotB, TempPar, VisCent, and VisPeri (all *p*s *<* 0.0015). SomMotA was less flexible than all systems except SomMotB, TempPar, and Amygdala (all *p*s *<* 0.0037).

The Limbic system had the second most flexible nodes after ContC (median flexibility = 0.845), and was more flexible than DefaultA, DefaultB, DorsAttnA, SomMotA, SomMotB, TempPar, VisCent, and VisPeri (all *p*s *<* 0.005). Subcortical nuclei varied in their flexibility, with median flexibility ranging from 0.845 in the Cerebellum to 0.722 in the Amygdala. Ultimately, none of the subcortical nuclei were statistically distinguishable from one another. The only coarse-grained system that had distinguishable fine-grained systems was Default Mode: DefaultC was more flexible than DefaultA (*p* = 0.004).

The relative lack of flexibility in somatomotor, temporoparietal, and visual systems suggests that these systems are largely indifferent to perceived social support. The systems with the most flexible nodes, on the other hand, (e.g. ContC and Limbic, and to a lesser extent Cerebellum and DefaultC) may show patterns of activity that are more sensitive to the differences between the seven perceived measures of social support. In other words, given the connectivity of these more flexible systems, the measures of perceived social support are more differentiable.

#### Flexibility across social support measures

Are there particular social support layers that are substantially different from the rest, forcing nodes to switch their community affiliation and therefore driving flexibility higher? To address this question, we looked at the proportion of nodes that switched their community affiliation between each pair of layers (see Figure 5c). To construct a null model, for each partition and for each node we permuted community affiliations across layers 1000 times. We then computed the proportion of nodes that changed communities between every pair of layers and took the average across partitions. To obtain a *p*-value, we divided this average by the number of permutations. The proportion of nodes that switched their communities between the *Friendship* layer and every other layer was significantly larger in the observed data than in the permuted data (all *p*s *<* 0.0001). No other layer had significantly different proportions of nodes changing for *every* comparison, but *Instrumental Support* had the next most. The proportions of nodes changing communities was significantly larger for *Instrumental Support* than all layers *except Hostility* (all *p*s *≤* 0.03). This suggests that communities, and by extension the roles nodes play, in the *Friendship* layer were especially different from all the other layers. Communities in the *Instrumental Support* layer also tended to be different from most other layers.

How similar are systems to this global pattern, in which the *Friendship* and *Instrumental Support* layers contain relatively distinct communities? To determine this, we computed the proportions of nodes changing communities between each pair of layers both (1) for the whole brain and (2) for each system. We then correlated these lists of proportions. All systems except Somatomotor and Temporoparietal (which are both generally inflexible; see Figure 5b) had significant correlations with the global pattern (all *r*s*≥*0.56, all *p*s *≤*0.0009; see Figure 5e). DorsAttn, Control, and Vis were the systems most similar to the global pattern (all *r*s *≥* 0.79); they were also most similar to each other (all *r*s *≥* 0.81; see Figure 5d), even though they varied greatly in the flexibility of their nodes (see Figure 5b). Temporoparietal, on the other hand, was dissimilar from all other systems (*r* with Default = 0.47, all other *r*s *≤* 0.22). Similarity to the global pattern of Friendship driving flexibility was not observed only in more flexible systems, like Control; less flexible systems, like Vis, also showed this pattern.

## Discussion

We set out to link different experiences in the social environment to variation in brain network organization. We use the established methods of community detection (Newman and Girvan, 2004) and multilayer modeling (Mucha et al., 2010) in multiply innovative ways to make this connection. We highlight two findings that contribute to research on the neural sensitivity to social support. First, the subcortex and control systems are especially sensitive to the different constructs of perceived social support. Network nodes in these systems are highly flexible; their community affiliations differ at high rates across social support measures. Nodes in the somatomotor, temporoparietal, and visual systems, on the other hand, were relatively stable in their community affiliations. Second, the *Friendship* layer, and to a lesser extent *Instrumental Support*, drove flexibility; higher proportions of nodes across all systems switched their community affiliations between *Friendship* (and *Instrumental Support*) and all other layers.

When examining the stability of our results across processing pipelines, we find that correlational patterns between brain network edge weights and scores on seven measures of social support are consistent across resting fMRI scans and with and without global signal regression. While many edges do not survive permutation testing, notably many in the subcortex do. We examine the parameter space of the multilayer model in detail according to variability in flexibility; according to similarity of partitions to each other, to consensus communities, and to canonical systems; and according to modes of variation based on PCA. In general there is more variation across the resolution parameter (*γ*), which tunes the size and number of communities, than the coupling parameter (*ω*),which tunes the strength of the relationship between layers, in line with previous network neuroscience studies that use multilayer models (Betzel et al., 2019).

We zoom into two points in parameter space: the parameter value pairs for which the magnitudes of the coefficients of the first and second principal components are maximized. The point for PC1 yield 14 total communities, while the point for PC2 yields 25 total communities. In both cases, Somatomotor, Default, and Visual nodes mostly have low flexibility while Control and Subcortex (and to a lesser extent Attention and Limbic) nodes have a wider distribution to their flexibility values.

When we broke down flexibility across the brain and canonical systems (Yeo et al., 2011), we found that ContC had the most flexible nodes while SomMotA had the least flexible. Greater flexibility, in our case, suggests that the role(s) of nodes in a system differentiate between the measures of social support. Lower flexibility has two possible interpretations: (1) the nodes of a system do not distinguish between measures of social support because their activity is not driven by or correlated with social support or (2) the nodes of a system do not distinguish between measures of social support because their activity is similarly correlated with each measure. Disambiguating these two explanations might involve cross-referencing flexibility with system-level activation in social tasks. Given that previous studies have linked loneliness, for example, with the Default network (Mwilambwe-Tshilobo et al., 2019; Spreng et al., 2020), it might be the case that interpretation 1 applies to inflexible Somatomotor and Visual nodes since we would expect these brain regions to have a weaker relationship with social support. Interpretation 2 might apply to inflexible Default nodes.

Just as we broke down flexibility across brain systems, we also broke down flexibility across the social support measures. We find that, for principal component 1, node roles (operationalized here as community affiliation) are substantially different in the *Friendship* layer. The same is true to a lesser extent for the *Instrumental Support* layer. This may be due in part to the fact that the *Friendship* layer has more communities than any other layer and the *Instrumental Support* layer has fewer communities than any other layer. This means that any node assigned to a community unique to the *Friendship* layer has to change its affiliation when compared with another layer.

Notably, our layer-based analysis does not yield any results showing different patterns between layers indicating “positive” social experiences (e.g. *Friendship, Instrumental Support, Emotional Support*) and “negative” social experiences (e.g. *Stress, Hostility, Rejection, Loneliness*). Several neuroecological studies have proposed different dimensions of environmental stressors (Sheridan and McLaughlin, 2014; McLaughlin and Sheridan, 2016; Rice et al., view), including deprivation versus threat and social versus economic versus neighborhood. The indifference of community changes to “positive” versus “negative” experiences suggests valence may not be a meaningful dimension for the brain. A more salient distinction, according to previous literature, may be between emotional support and instrumental support (Morelli et al., 2015). While both measures capture aspects of social support, emotional support speaks more to empathy and instrumental support more to resource provision.

While the importance of control and subcortical nodes for social processes aligns with work in social neuroscience and neuroecology (Coan et al., 2017; Steinberg, 2017; Xin and Lei, 2015; Cacioppo et al., 2007), our approach provides rich detail that traditional region of interest analyses cannot. In particular, multilayer modeling allows us to take all social support variables into consideration, get detailed information about all nodes in the system, and make direct comparisons about nodes and systems across social support variables. This detailed information can be connected to canonical brain systems and social support, making the multilayer approach a bridge between subjective social measures and brain functional connectivity. To clarify what exactly the role of control and subcortical nodes might be, it is important to consider how the observed patterns of flexibility change across individuals according to their experiences in the social environment. That is, how do patterns of flexibility for control and subcortical nodes differ between individuals with consistent high quality social support and individuals who experience more rejection, hostility, loneliness, and stress in their social environment?

Our method offers innovation on commonplace network neuroscientific tools (Bassett et al., 2011; Vaiana and Muldoon, 2020) to provide details about brain network connectivity relationships not yet observed in neuroecological work. Moreover, this methodological innovation can be applied to *any* set of measures, whether they are cognitive, behavioral, or demographic. We want to highlight that our layer-based analysis (which identified that more nodes change community affiliation between *Friendship* and all other layers) is only feasible when there is all-to-all coupling between layers. In other words, applications of multilayer modeling to study time-varying functional connectivity cannot identify layer differentiation, because layers are often only coupled to adjacent layers. This layer-based analysis is thus a methodological innovation that helps shed light on how the role of brain systems might vary according to (in our case, social) context.

It is worth noting that our particular multilayer implementation, and by extension our layer-based analysis, can be applied to any set of interrelated measures. Our application of multilayer modeling to perceived social support is motivated by work in neuroecology (Gonzalez et al., 2015; Coan et al., 2017; Pelletier-Baldelli et al., 2022; DeCross et al., 2022; Morawetz et al., 2020), but the implementation’s agnosticism toward the measures used speaks to its generality: it is a powerful tool for a wide variety of research questions. This approach can be used to investigate how brain connectivity varies across experiences in the physical or economic environment or across different behavioral or cognitive measures.

## Limitations and future directions

Within a developmental neuroecological framework, variation across individuals is meaningful and can point to important differences in environmental experience. Critically, this study does not investigate individual or even group differences. By using group-averaged functional connectivity, we cannot speak directly to adaptations, individual differences, or precise neuroendophenotypes. Before such an approach can be taken, it was imperative to first show that network tools could even describe neuroendophenotypes. That our results align with neuroecological findings (e.g. Gonzalez et al., 2019, 2015; Coan et al., 2017) suggests this is the case. Therefore, future studies should explore how individual variation in brain networks relates to variation in experiences in the environment.

Because the HCP dataset includes only adults, we were limited in the developmental conclusions we can draw. Bigger effect sizes would be expected in adolescent populations in particular, since adolescent represents a critical period of rapid and high-stakes adaptation to the (especially social) environment (Rudolph et al., 2021; Gonzalez et al., 2015; Pelletier-Baldelli et al., 2022; DeCross et al., 2022). Applying methods similar to ours to a dataset like that of Adolescent Brain Cognitive Development study may be particularly illuminating.

## Conclusions

The present study offers theoretical and methodological innovations, both of which establish a link between the fields of neuroecology and network neuroscience. Our theoretical contribution is motivated by neuroecology. Neuroecological perspectives hold that individual differences in brain features are meaningful and are often related to differences in experience in the environment over the course of development (Gottlieb, 1991). While there are myriad environmental measures that could be linked to brains, we focus on the social environment. Although network neuroscientists have explored links between the social environment and brain networks before (Chan et al., 2018; Ellwood-Lowe et al., 2021; Mwilambwe-Tshilobo et al., 2019; Spreng et al., 2020; Krendl and Betzel, 2022; Barrett and Satpute, 2013; Noonan et al., 2018; Cassidy et al., 2021; Morawetz et al., 2020, 2021), the relationship between social support specifically and brain networks has been heretofore untouched. Our study addresses this gap explicitly and establishes a theoretical connection between neuroecology and network neuroscience by identifying how brain networks change in relation to social contexts.

Our methodological contribution is grounded in network science. Studying the relationship between social support and brain networks involves integrating rich environmental data with whole-brain data. In a way, this is structurally identical to the many brain-behavior association studies. We diverge from this resemblance by leveraging tools from network science to address neuroecological questions. We employ multilayer modeling to identify stability and variation in network community structure across seven measures of perceived social support. Because we construct the multilayer model to have all-to-all coupling, we are able to make pairwise comparisons between all layers to see how brain-environment relationships change across social support contexts. This methodology and our results align with and build on existing work in both network neuroscience (Bassett et al., 2011; Vaiana and Muldoon, 2020) and social neuroecology (Coan et al., 2017; Gonzalez et al., 2019), facilitating a rich connection between the two fields.

There are two key takeaways from this work: (1) dimensions of social environment are associated with variation in brain networks and (2) the methods used in this paper can be used to understand the interplay between brain and environment, behavior, cognition, or disease. This work serves as a foundation for future research at the intersection of network neuroscience and social neuroecology, or other fields.

## Materials and Methods

### Data

Our data included self-report social support data and resting state fMRI (rsfMRI) data from the Human Connectome Project (HCP) (Van Essen et al., 2013). For the social data, we used the seven perceived social support measures from the NIH Toolbox included in HCP data: *Friendship, Emotional Support, Instrumental Support, Hostility, Rejection, Loneliness, and Stress* (Cyranowski et al., 2013). All measures had eight items except *Loneliness*, which had five items. All items from all measures were rated on a five-point scale (ranging from “Never” to “Always”) and T-scored to fall between a range of 0 to 100, with 50 representing the population average. Within the present sample, the average score for each measure was: 51.11 *±* 9.38 for *Friendship*; 50.39 *±* 8.58 for *Emotional Support*, 49.79*±* 8.27 for *Instrumental Support*, 47.74*±*8.27 for *Hostility*, 51.91*±*9.66 for *Rejection*, 47.86*±*8.38 for *Loneliness*, and 47.76*±* 7.87 for *Stress* (see Supplemental Figures S1a,b for score distributions and correlations between scores). Social measures were typically assessed on the same day as or within one day of rs-fMRI data collection.

The HCP dataset (Van Essen et al., 2013) consisted of resting state functional magnetic resonance imaging (fMRI) scans (henceforth REST1 and REST2) from 100 unrelated adult subjects. These subjects were selected as they comprised the “100 Unrelated Subjects” released by the Human Connectome Project. After excluding subjects based on data completeness and quality control (see **Quality Control**), the final subset utilized included 95 subjects (56% female, mean age = 29.29 *±* 3.66, age range = 22-36, but see Supplemental Figure S2b for data about how results generalize with different sample sizes)). The study was approved by the Washington University Institutional Review Board and informed consent was obtained from all subjects. A comprehensive description of the imaging parameters and image preprocessing can be found in Glasser et al. (2013). Images were collected on a 3T Siemens Connectome Skyra with a 32-channel head coil. Subjects underwent two T1-weighted structural scans, which were averaged for each subject (TR = 2400 ms, TE = 2.14 ms, flip angle = 8^*°*^, 0.7 mm isotropic voxel resolution). Subjects underwent four resting state fMRI scans over a two-day span, henceforth we refer to the rsfMRI scans from day 1 as REST1 and the rsfMRI scans from day 2 as REST2. (See Supplemental Figure S2c for comparisons of our findings across REST1 and REST2.) The fMRI data was acquired with a gradient-echo planar imaging sequence (TR = 720 ms, TE = 33.1 ms, flip angle = 52^*°*^, 2 mm isotropic voxel resolution, multiband factor = 8). Each resting state run duration was 14:33 min, with eyes open and instructions to fixate on a cross.

### Quality Control

All preprocessed time series were visually inspected for visual artifact. Subject motion measurements during the fMRI scanning sessions were obtained from the HCP minimal preprocessing pipeline output directories (files: `Movement RelativeRMS.txt` and `eddy unwarped images.eddy movement rms`).

Across fMRI sessions and within an fMRI session, the mean and mean absolute deviation of the motion measurements were calculated, resulting in four summary motion measures per subject. Subjects without complete data or exceeding 1.5 times the inter-quartile range (in the adverse direction) of the measurement distribution for more than one of these summary measurements were excluded. We discarded frames *>* 0.15 motion threshold. After these quality assurance steps, data from 92 subjects remained.

### Image Processing

Functional images of the HCP dataset were minimally preprocessed according to the description provided in Glasser et al. (2013). Briefly, T1w images were aligned to MNI space before undergoing FreeSurfer’s (version 5.3) cortical reconstruction workflow. fMRI images were corrected for gradient distortion, susceptibility distortion, and motion, and then aligned to the corresponding T1w with one spline interpolation step. This volume was further corrected for intensity bias and normalized to a mean of 10000. This volume was then projected to the *32k fs LR* mesh, excluding outliers, and aligned to a common space using a multi-modal surface registration (Robinson et al., 2014). The resultant CIFTI file for each HCP subject used in this study followed the file naming pattern:

`* REST`*{*`1,2`*} {*`LR,RL`*}* `Atlas MSMAll.dtseries.nii`.

### Parcellation

As HCP fMRI was provided in *32k fs LR* space, this data could be parcellated based on the available Schaefer 400 parcellation (Schaefer et al., 2018) in the CIFTI file format. Additionally, a novel gradient-based subcortical parcellation was used to delineate nodes within the amygdala, hippocampus, thalamus, and striatum consisting of 41 regions per hemisphere (Scale III parcellation from Tian et al., 2020). Volumetric ROIs of subcortical and cerebellar (Diedrichsen et al., 2009) regions were defined in the MNI152 1mm space, conforming to the cifti specification. We also compared coarser 200 cortical node and 100 cortical parcellations to this maximally fine-grained parcellation (see Supplemental Figure 2c).

### Functional Connectivity

Each preprocessed BOLD image was linearly detrended, band-pass filtered (0.008-0.08 Hz), confound regressed and standardized using Nilearn’s `signal.clean` function, which removes confounds orthogonally to the temporal filters. The confound regression strategy included six motion estimates, mean signal from a white matter, cerebrospinal fluid, and whole brain mask, derivatives of these previous nine regressors, and squares of these 18 terms. Spike regressors were not applied to the HCP data. The 36 parameter strategy (with and without spike regression) has been show to be a relatively effective option to reduce motion-related artifacts (Parkes et al., 2018). Following these preprocessing operations, the mean signal was taken at each node in the surface space. We report on global signal regressed data in the main text but provide comparisons to data without global signal regression in Supplemental Figure S1a.

### Correlating functional connectivity and social support

For each resting scan, we computed functional connectivity using the Pearson correlation between all 482 nodes for each individual see Figure 1a - c). This gave us 92 matrices of dimension 482 by 482, whose entries *ij* represent the subject-specific edge weight between nodes *i* and *j* (see Figure 1a). We then vectorized these subject-specific functional connectivity matrices to obtain a [1*×*115921] vector of edge weights. Subjects’ edge weight vectors were concatenated to create a [92 *×* 115921] edge weight matrix (see Figure 1c. This allowed us to calculate the Spearman correlation between scores on each of the seven social support measures (see Figure 1b) and all edge weights, resulting in a [7 *×* 115921] matrix.

Each of these vectorized correlations could be returned to the upper triangle of a [482*×* 482] matrix, resulting in seven such matrices, where the element *ij* of matrix *k* gave the correlation between the edge weight of nodes *i* and *j* and social support measure *k* (see Figure 1d). These seven correlation matrices had a high correlation across REST1 and REST2 (see Supplemental Figure S1c), with and without global signal regression (see Supplemental Figure S2a), across sample sizes (see Supplemental Figure S2b, and across parcellations of cortex into 100, 200, and 400 regions (see Supplemental Figure S2c. The results of those analyses are reported in Supplemental Figure S1. The results presented in the remaining sections focus on correlation values averaged over both REST1 an REST2 scans based on data processed using a pipeline that includes global signal regression and a cortical parcellation of 400 nodes with 82 subcortical parcels.

### Examining brain-behavior correlation patterns

Rather than examine the correlation patterns independently for each social support measure, we study them jointly, representing each as a layer in a multi-layer network. This approach has been used previously Bassett et al. (2011, 2013); Finc et al. (2020); Betzel et al. (2019), though to the the best of our knowledge this is the first time that the layers have corresponded to patterns of brain-behavior correlations, as opposed to functional connectivity. Multilayer networks provide a framework to study different versions of a network at once. In a multilayer model, each layer is an individual network (e.g. a single functional connectivity network) which can be linked to other layers to capture change in a network over time or in different contexts. This allows any network computation to be performed on all layers simultaneously, providing insight into how a network changes over time or across different contexts (Mucha et al., 2010).

We sought to quantify which patterns were consistent and which fluctuated across the social support measures. To do this we used community detection. If we were to identify the community structure for each social support measure individually, there would be no principled way to compare structure across social support measures. We cannot know whether Community 1 for one measure is referring to the same set of nodes as Community 1 for another measure if we perform community detection on the layers independently, because the community detection algorithm involves some stochasticity. Furthermore, because the seven social support measures are strongly correlated with one another–some positively and some negatively–we want to take all measures into account when performing any analysis. Not doing so would provide redundant information. To accommodate this, we used multilayer modularity maximization (Newman and Girvan, 2004). Modularity maximization (both the single- and multi-layer versions) aims to identify groups of nodes who connectivity to one another exceeds that of a chance model. Here, however, we aim to detect groups of brain regions with similar brain-behavior correlation profiles. We use a uniform null model and the Louvain algorithm to maximize the modularity function *Q*:

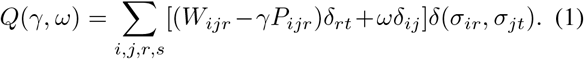

Nodes are linked to each other across layers through the resolution parameter *ω*. The value of *ω* influences the homogeneity of communities across layers (indicated by *r* and *t*), such that small *ω* values emphasize layer-specific communities and large *ω* values identify communities shared across layers. *W*_*i*_*j* and *P*_*ij*_ are the actual and expected weights of the link connecting nodes *i* and *j. s∈* 1, …, *K* indicates to which cluster node *i* belongs. The Kronecker delta function, *δ*(*x, y*) takes a value of 1 when *x* = *y* and a value of 0 otherwise. The parameter *γ* represents a spatial resolution weight that scales the influence of the null model. The optimization of *Q*(*γ, ω*) returns for each layer a partition of the network into assortative communities (see Figure 1g).

#### Multilayer model

Before we applied multilayer modularity maximization to the multilayer model, a transformation of the data was necessary. When a network includes negative weights, as is the case with correlation-based networks, modularity maximization will treat all negatively weighted items as part of one community. Since this could misrepresent patterns of correlations found in our data, we transform the seven correlation matrices into similarity matrices. We do this by computing the similarity between every pair of rows in a given correlation matrix, such that the entry *ij* corresponded to the Pearson correlation between the correlation profiles of regions *i* and *j* (see Figure 1e). Each of the seven similarity matrices becomes itself a layer in the multilayer model, such that within a given layer we are given the relationships between scores on one social support measure and all edge weights for all subjects (see Figure 1f). By using a multilayer approach, we can run community detection on all social support layers with varying amounts of coupling between layers and track which nodes change their community affiliation between which layers (see Figure 1g). This particular formalization represents an innovation in applications of multilayer networks to neuroscience.

To build the multilayer model we flatten our seven similarity matrices into a 2-D tensor, such that each matrix (which is itself the similarity matrix of correlations between a social support measure and functional edge weights) is positioned on the diagonal of the tensor matrix (see Figure 1f). Off-diagonal entries represent the coupling parameter *ω*. If *ω* is 0, each layer is independent in any calculation performed. As *ω* approaches 1, all the layers are increasingly treated as a single object where the individual layers are indistinguishable. By varying *ω*, then, we see how the community structure of the individual layers differs from each other and from the community structure of the layers taken as a whole. We tested a range of nine logarithmically spaced *ω*s from 0.0001 to 1.

#### Community detection

Since we are interested in the community structure of this multilayer model, we ran modularity maximization (Newman and Girvan, 2004) with a uniform null model and the Louvain algorithm to maximize the modularity function *Q*. By tuning the resolution parameter *γ* we see differences in communities according to how large their average correlations are. Larger *γ*s result in communities for which the average correlation between members is high, yielding numerous small communities. Smaller *γ*s give communities for which the average correlation between members is low, yielding fewer and bigger communities. We tested a range of 19 linearly spaced *γ*s from 0 to 0.9. Using 100 iterations of the modularity maximization algorithm, we also computed consensus communities (see Figure 4a).

### Flexibility

Using the community assignments resulting from the community detection algorithm, we can calculate how much a given node changes its community affiliation between layers, or the flexibility *f*_*i*_ of node *i*:

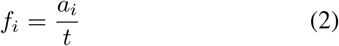

where *a*_*i*_ is the number of times node *i* changes its community affiliation and *t* is the total number of changes possible (see Figure 1h). When the coupling parameter *ω* is large, in general most nodes will have lower flexibility because the layers are so highly coupled that the amount a node can change is constrained. Similarly, when *ω* is small, most nodes will have higher flexibility.

---

**Figure S1:**
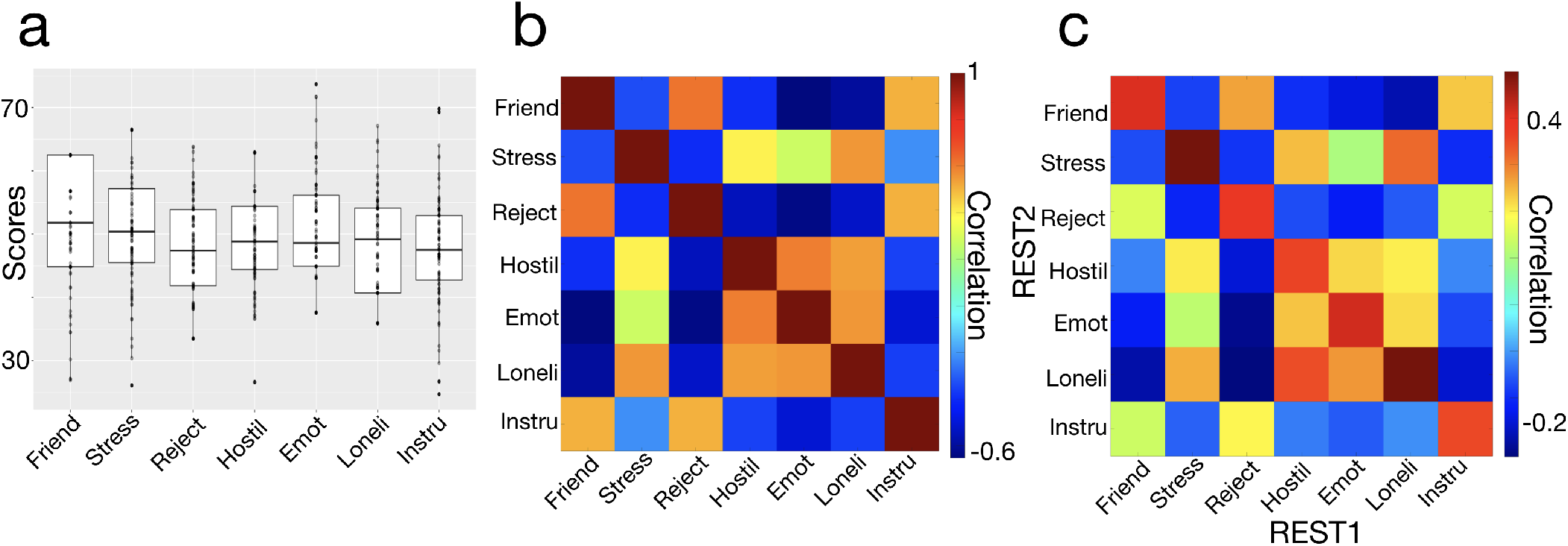
Social support measures. Panel (a) shows the distributions of scores for each of the social support measures. Panel (b) shows the correlations between all of the measures. Panel (c) shows the correlations across rest scans between each measure and the functional connectivity edge weights.

**Figure S2:**
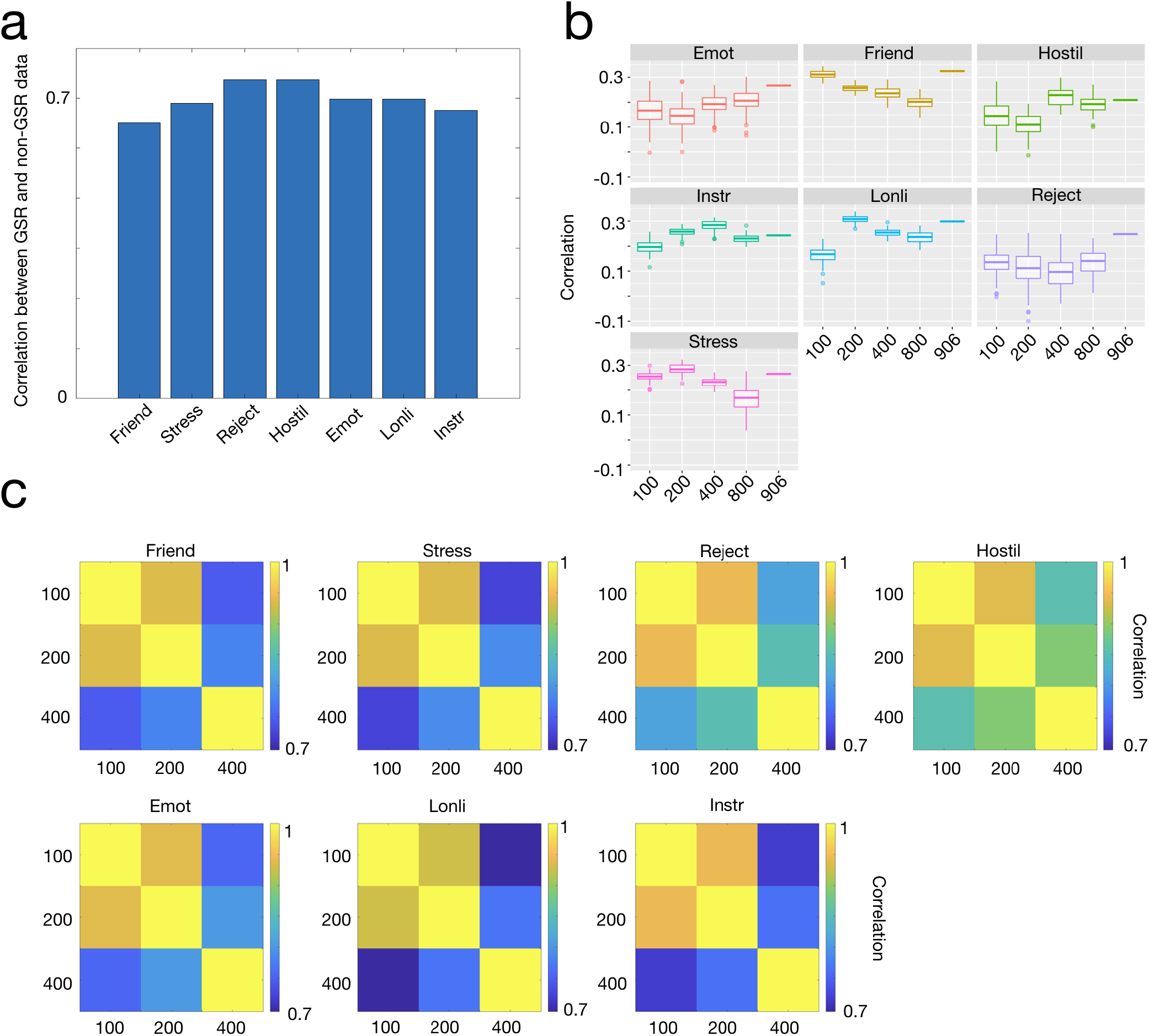
Generalizability of results. Panel (a) shows the correlations between data with and without global signal regression. For each measure, we correlate the matrices of correlations between edge weights and the social support measure. Correlations range from 0.551 (for the Friendship measure) to 0.6375 (for the Hostility measure). We also compared our original data to samples of various sizes from the HCP dataset (b). For each sample size (*n* = 100, 200, 400, 800), we subsampled *n* individuals at random from the entire HCP dataset. For each subsample, we computed the correlation between the original edge weight-social support correlation matrix and the subsampled edge weight-social support correlation matrix. Additionally, for all HCP subjects that had complete data (*n* = 906), we computed the correlation with the original data. All correlations are shown by measure in the box plots in panel (b). In panel (c), we show the correlations between the edge weight-social support correlation matrices across different parcellations. While the 100 node and 200 node parcellations tend to be more similar to each other than either is to the 400 node parcellation, all correlations are greater than 0.6968 (observed for the Loneliness measure).

**Figure S3:**
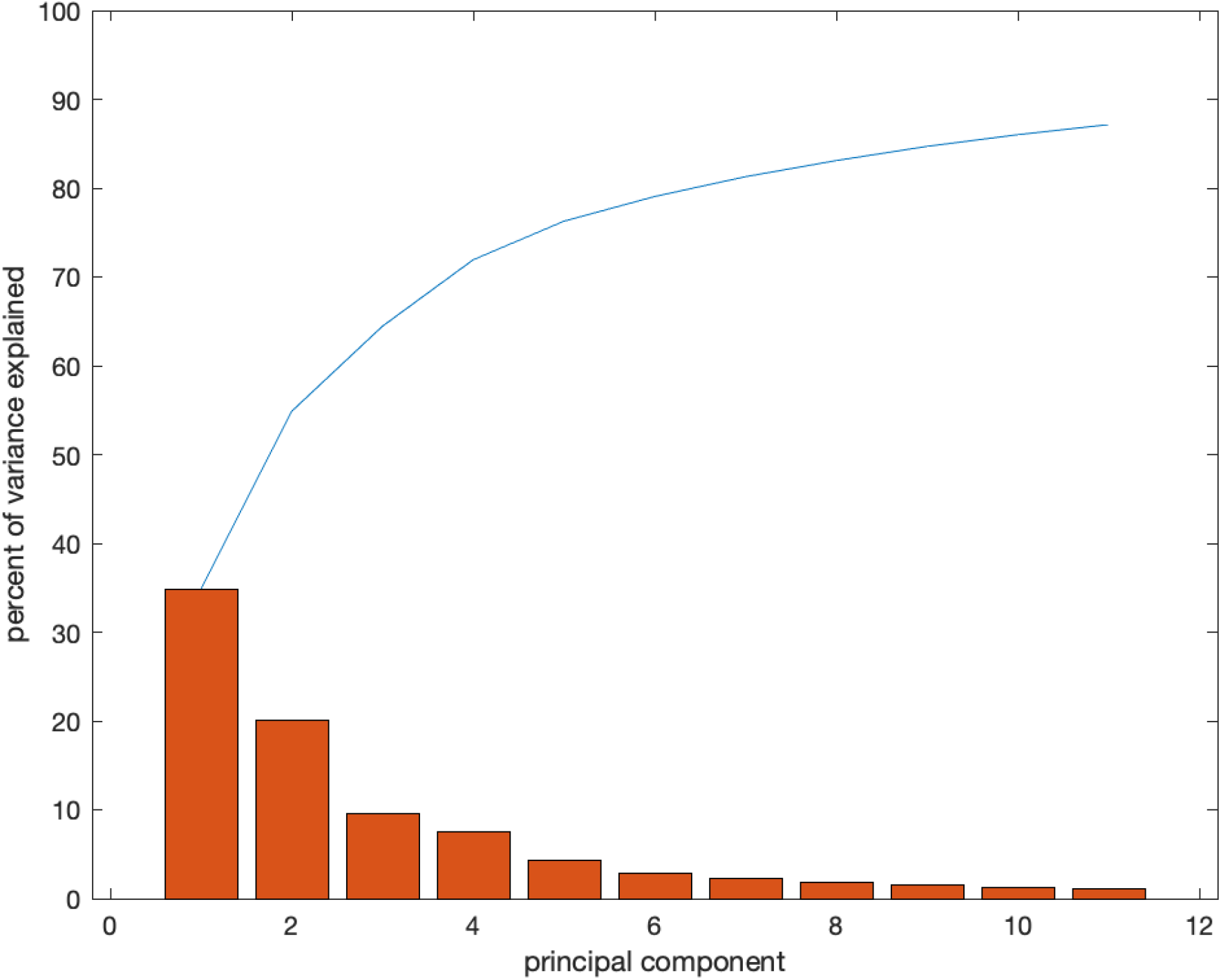
Cumulative variance explained. This figure shows the cumulative variance explained by the top 11 components that explain at least 1% of the total variance of parameter space each. Principal component 1 explains about 35% of the variance, principal component 2 explains just over 20% of the variance, and principal component 3 explains about 9.5%.

**Figure S4:**
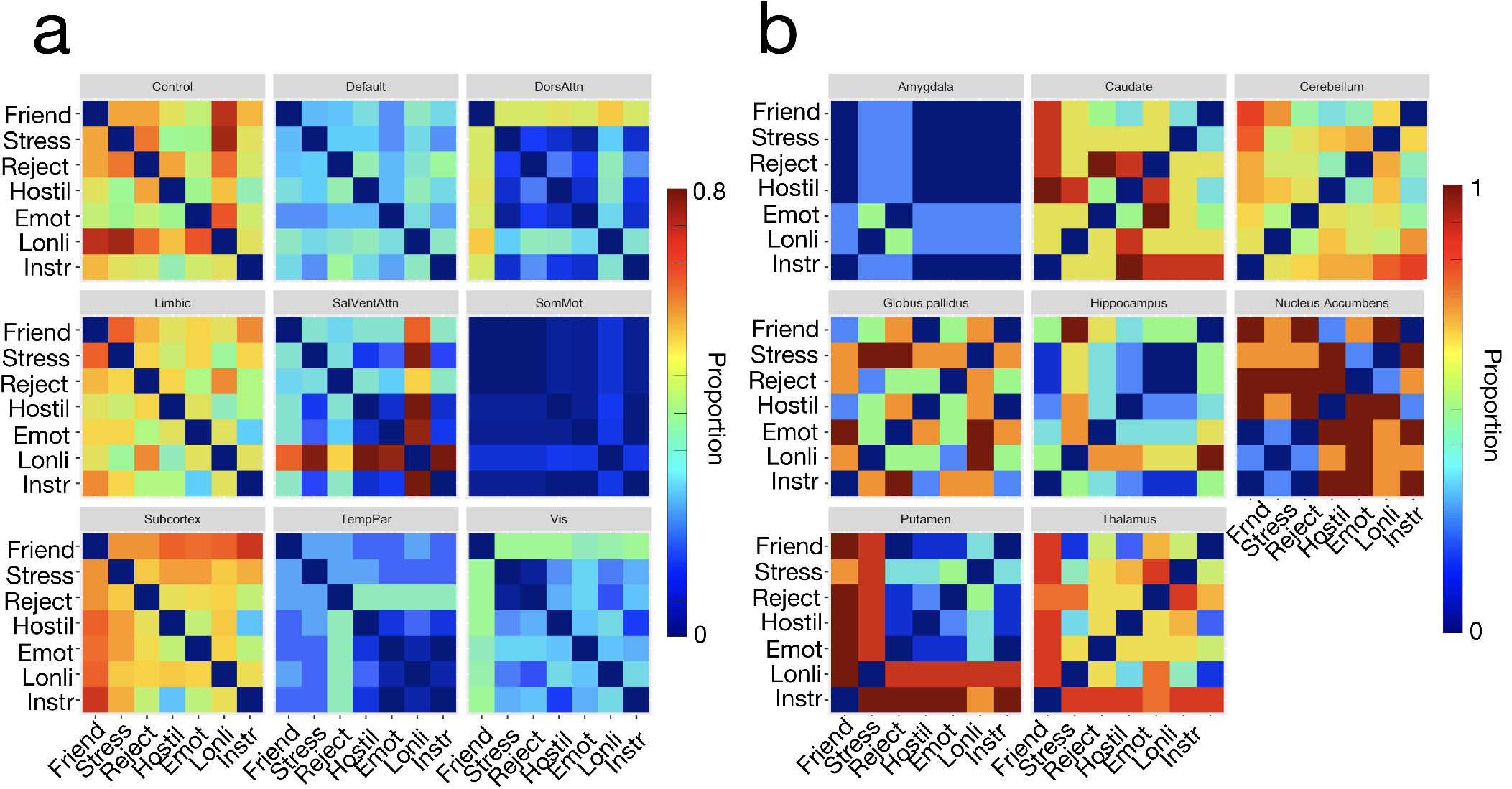
Breakdown of community switches between layers by system. In Figure 5 in the main text, we show the global pattern of the proportion of nodes that switch their community affiliation between each pair of layers. Here we break down for different canonical systems in panel (a) and for different subcortical nuclei in panel (b).

**Figure S5:**
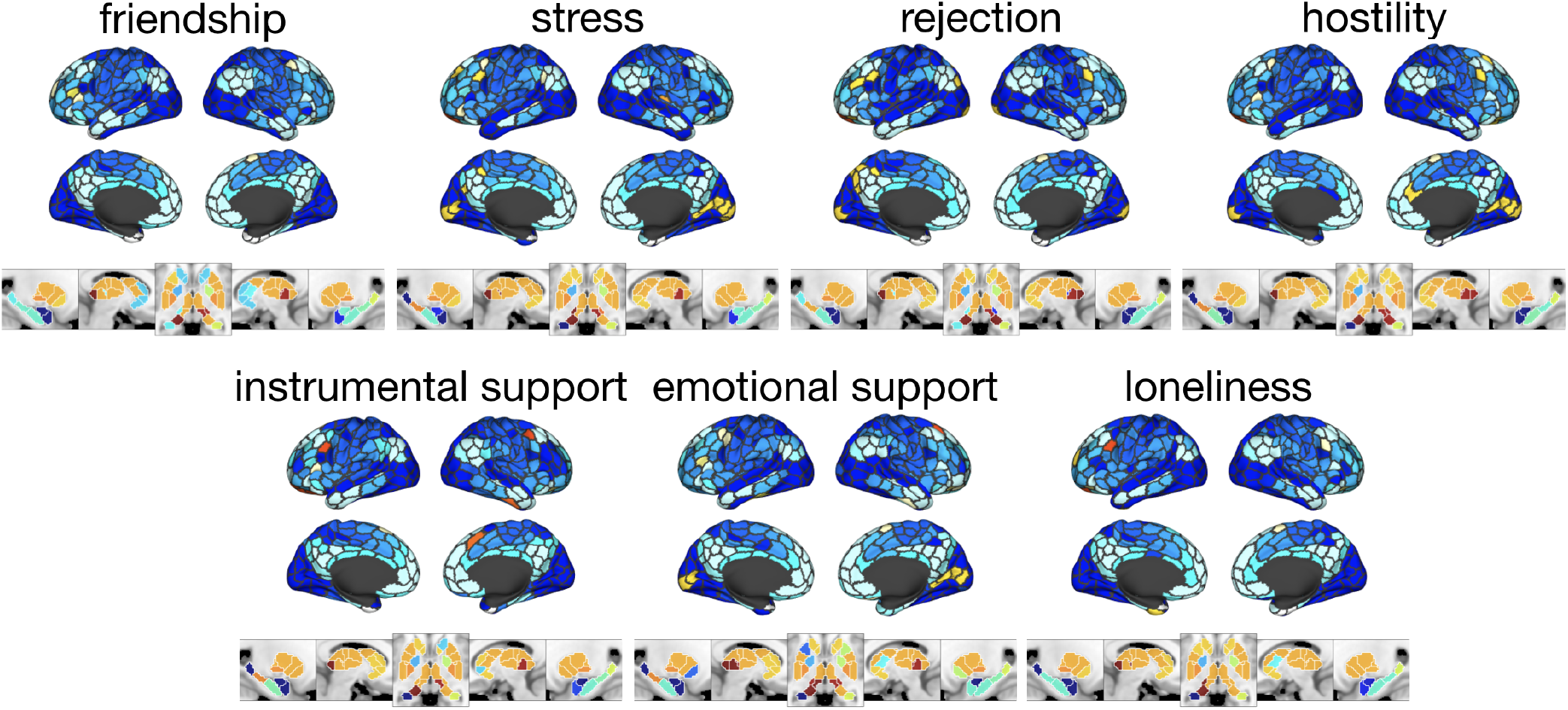
Communities for principal component 2. Community structure across all social support measures at the point in parameter space whose coefficient for principal component 2 has the largest magnitude (*γ* = 0.35, *ω* = 0.01). There are 25 communities total, all of which are present in each layer. The MNI coordinates for the five panels of subcortical communities are, from left to right: x = -23, x = 10, z = -3, x = 13, x = 27.

**Figure S6:**
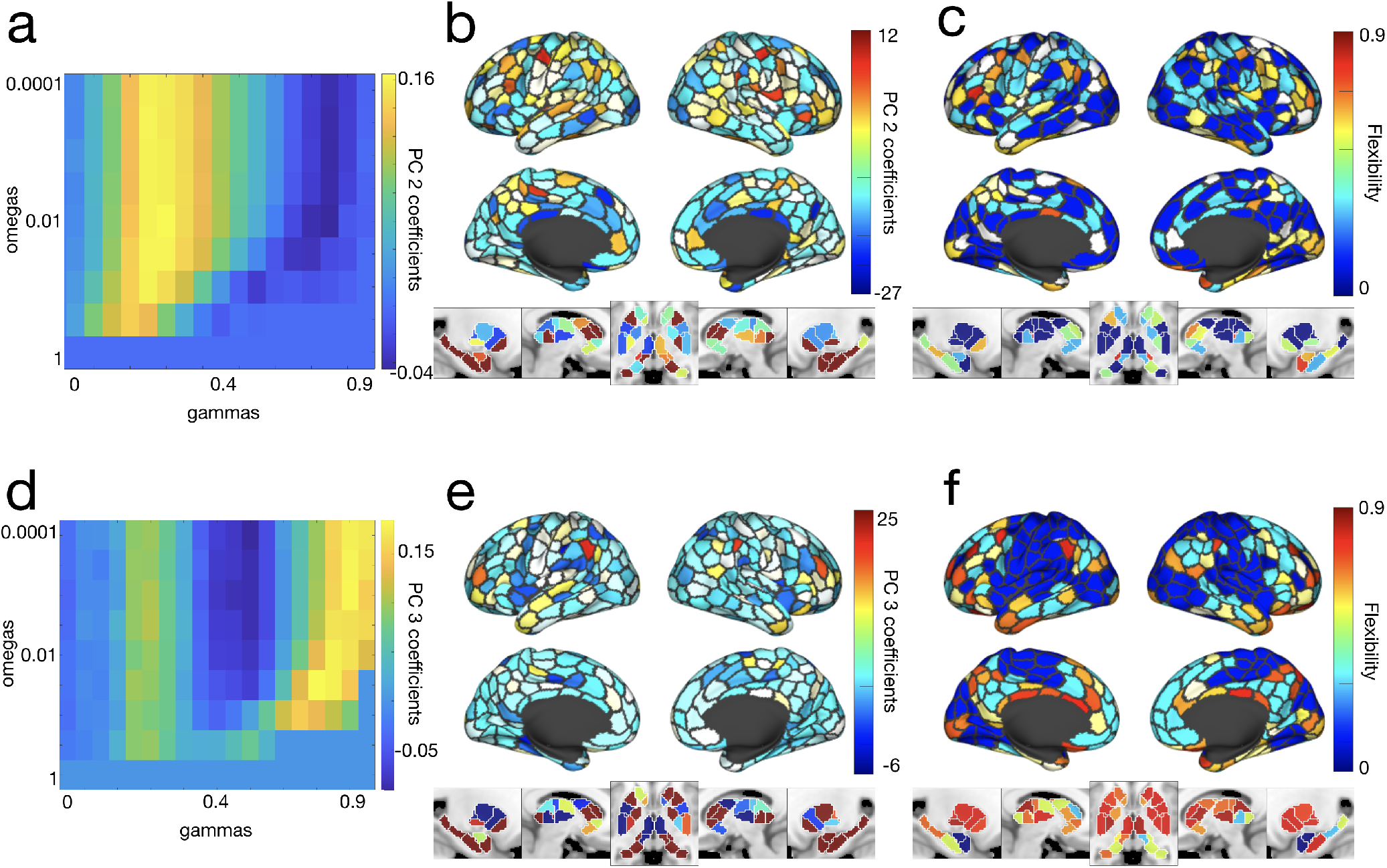
Principal components 2 and 3. The top row presents data from the second principal component, while the bottom row presents data from the third. Panels (a) and (d) show the PC coefficients for each point in parameter space. Panels (b) and (e) show the PC components projected into brain space. Panels (c) and (f) show the flexibility of cortical and subcortical nodes at the points in parameter space for which the magnitude of the coefficient of the PC is maximized. The MNI coordinates for the five panels of subcortical PC components and flexibility are, from left to right: x = -23, x = 10, z = -3, x = 13, x = 27.

## References

Barrett, L. F. and Satpute, A. B. (2013). Large-scale brain networks in affective and social neuroscience: towards an integrative functional architecture of the brain. Current opinion in neurobiology, 23(3):361–372.

Bassett, D. S., Porter, M. A., Wymbs, N. F., Grafton, S. T., Carlson, J. M., and Mucha, P. J. (2013). Robust detection of dynamic community structure in networks. Chaos: An Interdisciplinary Journal of Nonlinear Science, 23(1):013142.

Bassett, D. S., Wymbs, N. F., Porter, M. A., Mucha, P. J., Carlson, J. M., and Grafton, S. T. (2011). Dynamic reconfiguration of human brain networks during learning. Proceedings of the National Academy of Sciences, 108(18):7641–7646.

Baum, G. L., Ciric, R., Roalf, D. R., Betzel, R. F., Moore, T. M., Shinohara, R. T., Kahn, A. E., Vandekar, S. N., Rupert, P. E., Quarmley, M., et al. (2017). Modular segregation of structural brain networks supports the development of executive function in youth. Current Biology, 27(11):1561–1572.

Beckes, L. and Sbarra, D. A. (2022). Social baseline theory: State of the science and new directions. Current Opinion in Psychology, 43:36–41.

Berkman, L. F. and Syme, S. L. (1979). Social networks, host resistance, and mortality: a nine-year follow-up study of alameda county residents. American journal of Epidemiology, 109(2):186–204.

Betzel, R. F., Bertolero, M. A., Gordon, E. M., Gratton, C., Dosenbach, N. U., and Bassett, D. S. (2019). The community structure of functional brain networks exhibits scale-specific patterns of inter-and intra-subject variability. Neuroimage, 202:115990.

Betzel, R. F., Byrge, L., He, Y., Goñi, J., Zuo, X.-N., and Sporns, O. (2014). Changes in structural and functional connectivity among resting-state networks across the human lifespan. Neuroimage, 102:345–357.

Betzel, R. F., Satterthwaite, T. D., Gold, J. I., and Bassett, D. S. (2017). Positive affect, surprise, and fatigue are correlates of network flexibility. Scientific Reports, 7(1):1–10.

Cacioppo, J. T., Amaral, D. G., Blanchard, J. J., Cameron, J. L., Carter, C. S., Crews, D., Fiske, S., Heatherton, T., Johnson, M. K., Kozak, M. J., et al. (2007). Social neuroscience: Progress and implications for mental health. Perspectives on Psychological Science, 2(2):99–123.

Callaghan, B. L. and Tottenham, N. (2016). The neuro-environmental loop of plasticity: A cross-species analysis of parental effects on emotion circuitry development following typical and adverse caregiving. Neuropsychopharmacology, 41(1):163–176.

Cassidy, B. S., Hughes, C., and Krendl, A. C. (2021). Age differences in neural activity related to mentalizing during person perception. Aging, Neuropsychology, and Cognition, 28(1):143–160.

Chan, M. Y., Na, J., Agres, P. F., Savalia, N. K., Park, D. C., and Wig, G. S. (2018). Socioeconomic status moderates age-related differences in the brain’s functional network organization and anatomy across the adult lifespan. Proceedings of the National Academy of Sciences, 115(22):E5144–E5153.

Chen, E., Langer, D. A., Raphaelson, Y. E., and Matthews, K. A. (2004). Socioeconomic status and health in adolescents: The role of stress interpretations. Child development, 75(4):1039–1052.

Chen, E. and Miller, G. E. (2007). Stress and inflammation in exacerbations of asthma. Brain, behavior, and immunity, 21(8):993–999.

Chen, Z., He, Y., and Yu, Y. (2016). Enhanced functional connectivity properties of human brains during in-situ nature experience. PeerJ, 4:e2210.

Coan, J. A., Beckes, L., Gonzalez, M. Z., Maresh, E. L., Brown, C. L., and Hasselmo, K. (2017). Relationship status and perceived support in the social regulation of neural responses to threat. Social Cognitive and Affective Neuroscience, 12(10):1574–1583.

Cohen, S. (1988). Psychosocial models of the role of social support in the etiology of physical disease. Health psychology, 7(3):269.

Cohen, S. (2004). Social relationships and health. American psychologist, 59(8):676.

Cohen, S. and Wills, T. A. (1985). Stress, social support, and the buffering hypothesis. Psychological bulletin, 98(2):310.

Cyranowski, J. M., Zill, N., Bode, R., Butt, Z., Kelly, M. A., Pilkonis, P. A., Salsman, J. M., and Cella, D. (2013). Assessing social support, companionship, and distress: National institute of health (nih) toolbox adult social relationship scales. Health Psychology, 32(3):293.

De Jaegher, H., Di Paolo, E., and Gallagher, S. (2010). Can social interaction constitute social cognition? Trends in cognitive sciences, 14(10):441–447.

DeCross, S. N., Sambrook, K. A., Sheridan, M. A., Tottenham, N., and McLaughlin, K. A. (2022). Dynamic alterations in neural networks supporting aversive learning in children exposed to trauma: Neural mechanisms underlying psychopathology. Biological Psychiatry, 91(7):667–675.

Diedrichsen, J., Balsters, J. H., Flavell, J., Cussans, E., and Ramnani, N. (2009). A probabilistic mr atlas of the human cerebellum. neuroimage, 46(1):39–46.

Dosenbach, N. U., Nardos, B., Cohen, A. L., Fair, D. A., Power, J. D., Church, J. A., Nelson, S. M., Wig, G. S., Vogel, A. C., Lessov-Schlaggar, C. N., et al. (2010). Prediction of individual brain maturity using fmri. Science, 329(5997):1358–1361.

Ellwood-Lowe, M. E., Whitfield-Gabrieli, S., and Bunge, S. A. (2021). Brain network coupling associated with cognitive performance varies as a function of a child’s environment in the abcd study. Nature Communications, 12(1):1–14.

Fair, D. A., Dosenbach, N. U., Church, J. A., Cohen, A. L., Brahmbhatt, S., Miezin, F. M., Barch, D. M., Raichle, M. E., Petersen, S. E., and Schlaggar, B. L. (2007). Development of distinct control networks through segregation and integration. Proceedings of the National Academy of Sciences, 104(33):13507–13512.

Finc, K., Bonna, K., He, X., Lydon-Staley, D. M., Kühn, S., Duch, W., and Bassett, D. S. (2020). Dynamic reconfiguration of functional brain networks during working memory training. Nature communications, 11(1):1–15.

Gee, D. G., Gabard-Durnam, L. J., Flannery, J., Goff, B., Humphreys, K. L., Telzer, E. H., Hare, T. A., Bookheimer, S. Y., and Tottenham, N. (2013). Early developmental emergence of human amygdala–prefrontal connectivity after maternal deprivation. Proceedings of the National Academy of Sciences, 110(39):15638–15643.

Glasser, M. F., Sotiropoulos, S. N., Wilson, J. A., Coalson, T. S., Fischl, B., Andersson, J. L., Xu, J., Jbabdi, S., Webster, M., Polimeni, J. R., et al. (2013). The minimal preprocessing pipelines for the human connectome project. Neuroimage, 80:105–124.

Gonzalez, M. Z., Beckes, L., Chango, J., Allen, J. P., and Coan, J. A. (2015). Adolescent neighborhood quality predicts adult dacc response to social exclusion. Social cognitive and affective neuroscience, 10(7):921–928.

Gonzalez, M. Z., Coppola, A. M., Allen, J. P., and Coan, J. A. (2021). Yielding to social presence as a bioenergetic strategy: Preliminary evidence using fmri. Current research in ecological and social psychology, 2:100010.

Gonzalez, M. Z., Puglia, M. H., Morris, J. P., and Connelly, J. J. (2019). Oxytocin receptor genotype and low economic privilege reverses ventral striatum-social anxiety association. Social neuroscience, 14(1):67–79.

Gottlieb, G. (1991). Experiential canalization of behavioral development: Theory. Developmental psychology, 27(1):4.

Gross, E. B. and Medina-DeVilliers, S. E. (2020). Cognitive processes unfold in a social context: a review and extension of social baseline theory. Frontiers in Psychology, 11:378.

Hawkley, L. C. and Cacioppo, J. T. (2007). Aging and loneliness: Downhill quickly? Current Directions in Psychological Science, 16(4):187–191.

House, J. S., Landis, K. R., and Umberson, D. (1988). Social relationships and health. Science, 241(4865):540–545.

Krendl, A. C. and Betzel, R. F. (2022). Social cognitive network neuroscience. Social Cognitive and Affective Neuroscience, 17(5):510–529.

McLaughlin, K. A. and Sheridan, M. A. (2016). Beyond cumulative risk: A dimensional approach to childhood adversity. Current directions in psychological science, 25(4):239–245.

McLaughlin, K. A., Sheridan, M. A., and Lambert, H. K. (2014). Childhood adversity and neural development: deprivation and threat as distinct dimensions of early experience. Neuroscience & Biobehavioral Reviews, 47:578–591.

Miller, G., Chen, E., and Cole, S. W. (2009). Health psychology: Developing biologically plausible models linking the social world and physical health. Annual review of psychology, 60:501–524.

Morawetz, C., Berboth, S., and Bode, S. (2021). With a little help from my friends: The effect of social proximity on emotion regulation-related brain activity. Neuroimage, 230:117817.

Morawetz, C., Riedel, M. C., Salo, T., Berboth, S., Eickhoff, S. B., Laird, A. R., and Kohn, N. (2020). Multiple large-scale neural networks underlying emotion regulation. Neuroscience & Biobehavioral Reviews, 116:382–395.

Morelli, S. A., Lee, I. A., Arnn, M. E., and Zaki, J. (2015). Emotional and instrumental support provision interact to predict well-being. Emotion, 15(4):484.

Mucha, P. J., Richardson, T., Macon, K., Porter, M. A., and Onnela, J.-P. (2010). Community structure in time-dependent, multiscale, and multiplex networks. Science, 328(5980):876–878.

Mwilambwe-Tshilobo, L., Ge, T., Chong, M., Ferguson, M. A., Misic, B., Burrow, A. L., Leahy, R. M., and Spreng, R. N. (2019). Loneliness and meaning in life are reflected in the intrinsic network architecture of the brain. Social cognitive and affective neuroscience, 14(4):423–433.

Newman, M. E. and Girvan, M. (2004). Finding and evaluating community structure in networks. Physical review E, 69(2):026113.

Noonan, M., Mars, R., Sallet, J., Dunbar, R., and Fellows, L. (2018). The structural and functional brain networks that support human social networks. Behavioural brain research, 355:12–23.

Parkes, L., Fulcher, B., Yücel, M., and Fornito, A. (2018). An evaluation of the efficacy, reliability, and sensitivity of motion correction strategies for resting-state functional mri. Neuroimage, 171:415–436.

Pelletier-Baldelli, A., Sheridan, M. A., Glier, S., Rodriguez-Thompson, A., Gates, K. M., Martin, S., Dichter, G. S., Patel, K. K., Bonar, A. S., Giletta, M., et al. (2022). Social goals in girls transitioning to adolescence: Associations with psychopathology and brain network connectivity. Social Cognitive and Affective Neuroscience.

Puxeddu, M. G., Faskowitz, J., Sporns, O., Astolfi, L., and Betzel, R. F. (2022). Multi-modal and multi-subject modular organization of human brain networks. bioRxiv.

Rakesh, D., Zalesky, A., and Whittle, S. (2021). Similar but distinct–effects of different socioeconomic indicators on resting state functional connectivity: Findings from the adolescent brain cognitive development (abcd) study®. Developmental cognitive neuroscience, 51:101005.

Repetti, R. L., Taylor, S. E., and Seeman, T. E. (2002). Risky families: family social environments and the mental and physical health of offspring. Psychological bulletin, 128(2):330.

Rice, M., Merritt, H., and Gonzalez, M. Z. (Under Review). Dimensional approach to els reveals unique contribution of social early life stress to motivational phenotypes in emerging adulthood. Current Research in Ecological and Social Psychology.

Robinson, E. C., Jbabdi, S., Glasser, M. F., Andersson, J., Burgess, G. C., Harms, M. P., Smith, S. M., Van Essen, D. C., and Jenkinson, M. (2014). Msm: a new flexible framework for multimodal surface matching. Neuroimage, 100:414–426.

Rudolph, K. D., Davis, M. M., Skymba, H. V., Modi, H. H., and Telzer, E. H. (2021). Social experience calibrates neural sensitivity to social feedback during adolescence: A functional connectivity approach. Developmental Cognitive Neuroscience, 47:100903.

Saxbe, D. E., Beckes, L., Stoycos, S. A., and Coan, J. A. (2020). Social allostasis and social allostatic load: A new model for research in social dynamics, stress, and health. Perspectives on Psychological Science, 15(2):469–482.

Schaefer, A., Kong, R., Gordon, E. M., Laumann, T. O., Zuo, X.-N., Holmes, A. J., Eickhoff, S. B., and Yeo, B. T. (2018). Local-global parcellation of the human cerebral cortex from intrinsic functional connectivity mri. Cerebral cortex, 28(9):3095–3114.

Schmälzle, R., Brook O’Donnell, M., Garcia, J. O., Cascio, C. N., Bayer, J., Bassett, D. S., Vettel, J. M., and Falk, E. B. (2017). Brain connectivity dynamics during social interaction reflect social network structure. Proceedings of the National Academy of Sciences, 114(20):5153–5158.

Seeman, T. E. and McEwen, B. S. (1996). Impact of social environment characteristics on neuroendocrine regulation. Psychosomatic medicine, 58(5):459–471.

Sheridan, M. A. and McLaughlin, K. A. (2014). Dimensions of early experience and neural development: deprivation and threat. Trends in cognitive sciences, 18(11):580–585.

Spreng, R. N., Dimas, E., Mwilambwe-Tshilobo, L., Dagher, A., Koellinger, P., Nave, G., Ong, A., Kernbach, J. M., Wiecki, T. V., Ge, T., et al. (2020). The default network of the human brain is associated with perceived social isolation. Nature communications, 11(1):1–11.

Steinberg, L. (2017). A social neuroscience perspective on adolescent risk-taking. In Biosocial Theories of Crime, pages 435–463. Routledge.

Tian, Y., Margulies, D. S., Breakspear, M., and Zalesky, A. (2020). Topographic organization of the human subcortex unveiled with functional connectivity gradients. Nature neuroscience, 23(11):1421–1432.

Uchino, B. N., Cacioppo, J. T., and Kiecolt-Glaser, J. K. (1996). The relationship between social support and physiological processes: a review with emphasis on underlying mechanisms and implications for health. Psychological bulletin, 119(3):488.

Uchino, B. N., Landvatter, J., Zee, K., and Bolger, N. (2020). Social support and antibody responses to vaccination: a meta-analysis. Annals of Behavioral Medicine, 54(8):567–574.

Uchino, B. N., Trettevik, R., Kent de Grey, R. G., Cronan, S., Hogan, J., and Baucom, B. R. (2018). Social support, social integration, and inflammatory cytokines: A meta-analysis. Health Psychology, 37(5):462.

Uddin, L. Q., Supekar, K. S., Ryali, S., and Menon, V. (2011). Dynamic reconfiguration of structural and functional connectivity across core neurocognitive brain networks with development. Journal of neuroscience, 31(50):18578–18589.

Vaiana, M. and Muldoon, S. F. (2020). Multilayer brain networks. Journal of Nonlinear Science, 30(5):2147–2169.

Van Essen, D. C., Smith, S. M., Barch, D. M., Behrens, T. E., Yacoub, E., Ugurbil, K., Consortium, W.-M. H., et al. (2013). The wu-minn human connectome project: an overview. Neuroimage, 80:62–79.

Williams, W. C., Morelli, S. A., Ong, D. C., and Zaki, J. (2018). Interpersonal emotion regulation: Implications for affiliation, perceived support, relationships, and well-being. Journal of personality and social psychology, 115(2):224.

Wymbs, N. F., Orr, C., Albaugh, M. D., Althoff, R. R., O’Loughlin, K., Holbrook, H., Garavan, H., Montalvo-Ortiz, J. L., Mostofsky, S., Hudziak, J., et al. (2020). Social supports moderate the effects of child adversity on neural correlates of threat processing. Child abuse & neglect, 102:104413.

Xiao, M., Chen, X., Yi, H., Luo, Y., Yan, Q., Feng, T., He, Q., Lei, X., Qiu, J., and Chen, H. (2021). Stronger functional network connectivity and social support buffer against negative affect during the covid-19 outbreak and after the pandemic peak. Neurobiology of stress, 15:100418.

Xin, F. and Lei, X. (2015). Competition between frontoparietal control and default networks supports social working memory and empathy. Social cognitive and affective neuroscience, 10(8):1144–1152.

Yeo, B. T., Krienen, F. M., Sepulcre, J., Sabuncu, M. R., Lashkari, D., Hollinshead, M., Roffman, J. L., Smoller, J. W., Zöllei, L., Polimeni, J. R., et al. (2011). The organization of the human cerebral cortex estimated by intrinsic functional connectivity. Journal of neurophysiology.

Zaki, J. and Williams, W. C. (2013). Interpersonal emotion regulation. Emotion, 13(5):803.

